# Temperature affects the repeatability of evolution in the microbial eukaryote *Tetrahymena thermophila*

**DOI:** 10.1101/2020.11.11.378919

**Authors:** Jason A Tarkington, Rebecca Zufall

## Abstract

Evolutionary biologists have long sought to understand what factors affect the repeatability of adaptive outcomes. To better understand the role of temperature in determining the repeatability of adaptive trajectories, we evolved populations of different genotypes of the ciliate *Tetrahymena thermophila* at low and high temperatures and followed changes in growth rate over 4,000 generations. As expected, growth rate increased with a decelerating rate for all populations; however, there were differences in the patterns of evolution at the two temperatures. The growth rates of the different genotypes converged as evolution proceeded at both temperatures, but this convergence was quicker at the higher temperature. Likewise, we found greater repeatability of evolution, in terms of change in growth rate, among replicates of the same genotype at the higher temperature. Finally, we found no evidence of trade-offs in fitness between temperatures, but did observe asymmetry in the correlated responses, whereby evolution in a high temperature increases growth rate at the lower temperature significantly more than the reverse. These results demonstrate the importance of temperature in determining the repeatability of evolutionary trajectories.

## Introduction

The evolutionary trajectories of both natural and experimental populations are often remarkably similar to each other (Lenski and Travisano 1994; Colosimo et al. 2005; Woods et al. 2006; Conte et al. 2012; Nosil et al. 2018). However, there can also be substantial differences in the trajectories of initially identical experimental (Blount et al. 2008) and natural populations (Dieckmann and Doebeli 1999; McKinnon and Rundle 2002; Barluenga et al. 2006). While these types of studies have provided valuable insights into the repeatability of evolutionary trajectories, we still lack a comprehensive understanding of what conditions are likely to constrain trajectories from diverging due to stochastic forces, and thus contribute to the repeatability of evolution.

Previous work has demonstrated that temperature can fundamentally alter evolutionary outcomes, for example by increasing biological diversity at lower latitudes (Roy et al. 2002; Gillooly et al. 2004; Allen et al. 2006). One purported explanation for the effect of temperature is that mutation rates are different at different temperatures. However, empirical results are mixed, with some results showing higher mutation rates at higher temperatures, others lower rates at higher temperatures, and yet others are inconclusive (Faberge and Beale 1942; Kiritani 1959; Lindgren 1972). The “hotter is better” hypothesis predicts that warm-adapted populations will have higher maximum performance than their cold-adapted counterparts because of the evolution of greater robustness due to the inherently higher rates of biochemical reactions at higher temperatures (Huey and Bennett 1987; Angilletta et al. 2010). Evidence from comparative and experimental populations largely supports this hypothesis (e.g., Knies et al. 2009), however, again, some results are mixed (reviewed in Angilletta et al. 2010). Evidence from lab evolved *Escherichia coli* shows that greater fitness gains occur at higher temperatures and that populations evolved at lower temperature show trade-offs at higher temperatures but not vice-versa (Bennett and Lenski 1993; Mongold et al. 1996). Later work suggested that while the genetic changes underlying temperature adaptation were temperature specific, these mutations were also beneficial across all temperatures (Deatherage et al. 2017), demonstrating that at least for the most relevant mutations the observed trade-offs are not due to antagonistic pleiotropy. While tradeoffs could still result from the cumulative effect of less impactful mutations that show antagonistic pleiotropy it is striking that the most impactful mutations do not show any antagonistic pleiotropy and suggests the trade-offs, in part, result from mutation accumulation at sites that are relevant at the alternative temperature but neutral at the evolution temperature. Overall these results demonstrate that temperature fundamentally affects adaptive outcomes, yet it remains unknown whether the temperature at which a population evolves will also affect the repeatability of adaptive trajectories.

To assess how temperature affects the repeatability of evolution, we performed a longterm evolution experiment using the microbial eukaryote *Tetrahymena thermophila. T. thermophila* is useful as a model system due to its complex life history and development, and its ease of growth and tractability in lab (Nanney 1974; Merriam and Bruns 1988; Prescott 1994). The short generation time and small cell size mean that large populations can be evolved over many generations in the lab, and population size and growth rate are easily monitored. In addition, in contrast to most other microbes in which experimental evolution is regularly performed, it has a complex life history and genome structure (Nanney 1974; Merriam and Bruns 1988), allowing us to test whether the general patterns found in other microbes also apply to ciliates.

*T. thermophila*, like all ciliates, is notable for its genome structure. Two types of nuclei are maintained in each cell. The germline micronucleus (MIC) is diploid and transcriptionally silent during growth and asexual reproduction, while the somatic macronucleus (MAC) is 45-ploid and transcriptionally active, meaning it gives rise to the phenotype of the cell (Merriam and Bruns 1988). Ciliates are facultatively sexual, mostly reproducing asexually, but occasionally undergoing conjugative sex with cells of a different mating type (Nanney 1974). In our experiment, populations contained a single mating type, effectively preventing sex. Thus, only mutations that occurred in the MAC were subject to selection and captured in our fitness assays.

Two features of the *T. thermophila* genome may potentially impact the patterns of adaptive evolution. First, the polyploid MAC divides by amitosis, a process that results in the random distribution of alleles among daughter cells. Unlike with division by mitosis, amitosis results in allelic variation among asexual progeny (Doerder et al. 1992), which generates higher levels of genetic variation and potentially increases the rate of evolution. Second, *Tetrahymena* has an exceptionally low base-substitution mutation rate (Long et al. 2016), which has the potential to slow the rate of adaptation. However, the deleterious mutation rate is comparable to other species (Long et al. 2013), so the potential effect of mutation rate is currently unclear.

In this study, we conducted a long-term evolution experiment to determine how temperature affects repeatability of evolution in a ciliate. We evolved populations of different genotypes of *T. thermophila* in two different temperatures and monitored the fitness trajectories of replicate populations. To assess the effects of temperature on the dynamics of evolutionary trajectories, we asked: 1) Does the temperature at which populations evolve affect the future convergence or continued divergence of initial historical differences between genotypes, 2) Does evolution temperature affect the repeatability of fitness trajectories, and 3) How temperature-specific are adaptations, i.e., are there trade-offs or other correlated responses between temperatures? We predict that temperature plays an important role in way that variation is generated and acted on by selection. Thus, we expect that temperature will affect both the rate at which populations converge and the repeatability of evolution. Given prior results on trade-offs, we predict the populations evolving at a lower temperature are more likely to experience trade-offs.

We find that populations that evolved at the higher temperature tended to have higher fitness gains than their colder-evolved counterparts. The higher evolution temperature also led to faster convergence among populations started from different genotypes, and less divergence among replicate populations of a single starting genotype, indicating that evolution at the higher temperature does indeed result in more repeatable evolution. Finally, we found no indication of trade-offs, but rather an asymmetry in the correlated responses, whereby evolution at the higher temperature increases fitness at the lower temperature more than the reverse, possibly indicating greater environmental specificity of adaptations at the lower temperature.

## Methods

### Summary

We evolved 12 populations each at both 24°C and 37°C. Each set of 12 populations consisted of four replicate populations of three initial genotypes: two independent natural isolates and a hybrid progeny of these two isolates. Throughout the course of 4,000 generations of evolution, we measured growth rate at both 24°C and 37°C for each population.

### Strains and initial cross

Natural isolates of *T. thermophila*, designated 19617-1 (Tetrahymena Stock Center ID SD03089) and 19625-2 (Doerder 2019), were thawed from frozen stocks, inoculated into 5.5 mL of the nutrient rich medium SSP (Gorovsky et al. 1975) in a 50 mL conical tube, and incubated at 30°C with mixing for two days. These cultures were maintained as the parental lines. Eight populations were established for each genotype in 10 mL cultures in SSP. Four of these were maintained at 24°C and four at 37°C. These populations were designated by genotype (19617-1 or 19625-2, herein referred to as A and B, respectively) - replicate (1-4) - and evolution temperature (24°C or 37°C), e.g. A-1-37.

To generate the hybrid genotype from these strains, a conical tube of each parental genotype was centrifuged and the supernatant was poured off before the cells were resuspended in 10 μM Tris buffer (Bruns and Brussard 1974). After mixing at 30°C in Tris for two days to starve the cells and induce sexual competence, 1 mL of each starved parental population and an additional 1 ml of 10 μM Tris buffer were added to one well in a six-well plate and placed back in the 30°C incubator. The next morning (~12 hours later) the plate was checked for pairs and put back in the incubator for an additional 4 hours to allow progression of conjugation. Individual mating pairs were isolated under a microscope using a 2 μL-micropipette and placed in 180 μL of SSP in one well of a 96-well plate. The plate was then incubated for 48 hours after which time a single cell was isolated from each well and re-cultured into 180 μL of fresh SSP in a new well. After another 48 hours at 30°C four individual cells were isolated from one of the wells, into new wells with SSP, one for each of the replicate populations, and incubated at 30 °C for 48 hours. Each of the four 180 μL cultures was then split in two with each half being added to a separate 50 mL conical tube containing 10 mL of SSP, one designated for evolution at 37°C and the other at 24°C. These eight cultures are the starting hybrid populations and are designated as A×B (19625×19617) – replicate (1-4) – evolution temperature.

This provided us with a total of 24 populations consisting of three genotypes, two parental and one hybrid, half of which were evolved at 24°C and half at 37°C with four replicate populations of each genotype per treatment.

### Transfer regime

Approximately 25,000 cells (~90 μL) from each 37°C culture and 60,000 cells (~1 mL) from each 24°C culture were transferred to 10 mL of fresh SSP daily. Transfer volumes were adjusted as needed to maintain the same starting culture density at each transfer. On average, the 37°C evolved populations achieved ~6.8 generations per day and the 24°C populations achieved ~3.5 generations per day. This means that 37°C evolved populations experienced a wider range of densities during growth (~2,500 cells/mL - ~275,000 cells/mL) than the 24°C evolved populations (~6,000 cells/ mL —60,000 cells/mL), starting with a lower density and ending at a higher density. We estimate the effective population size to be approximately 100,000 cells for each evolved environment by calculating the harmonic mean of the population size at each discrete generation (Karlin 1968). To date, the 37°C populations have undergone ~9,000 generations of evolution and the 24°C populations have undergone ~4,000 generations of evolution. Here we describe the changes in growth rate over the first 4,000 generations of evolution at each temperature.

### Growth curves and analysis

As evolution progressed, growth rates of each population were measured at both 37°C and at 24°C, i.e. at both the temperature at which they evolved and the alternate temperature, on average every ~10-30 generations. Variation in number of generations between measurements arose because we could not perform 37°C and 24°C assays on the same days and the assays took different lengths of time at each temperature, thus we would do two consecutive single days of 37°C assays, followed by a single 24°C assay that lasted 2 days. Growth rate was measured by inoculating ~500 – 1000 cells into one well of a 96-well plate and measuring the optical density (OD) at 650 nm in a micro-plate reader every 5 minutes over the course of 24 – 48 hours for 37°C assays and 48 – 72 hours for 24°C assays (see below for validation of use of OD650 as a proxy for cell density). The maximum growth rate was then estimated for each well by fitting a linear regression to the steepest part of the growth curve (with OD on a log scale), estimating the maximum doublings per hour (h^-1^) (Wang et al. 2012; Long et al. 2013). 3 – 4 replicates of all populations were measured on a plate at each time point. ~375 plates containing 37°C evolved populations and ~625 plates containing 24°C evolved populations were run providing approximately 500 – 1,000 growth curves at either temperature per population over the 4,000 generations analyzed here.

### Validation of optical density as proxy for cell density

To validate that OD accurately measures cell density over a range of densities, cells from cultures growing on the micro-plate reader were counted under the microscope at several points during the growth cycle. 3-4 replicate wells were inoculated and the plate was run on the micro-plate reader at 37 °C. Every two to three hours, 5 μL of culture was removed and at least 200 cells were counted to estimate cell density. The cells were diluted as needed and then counted in 10 μL droplets containing approximately 40 cells. This process was independently repeated two times. The cell density measured by counting was tested for correlation with the OD measured by the micro-plate reader at each time point, and OD was found to be a good indicator of cell density (Pearson’s correlation coefficient = 0.9602; Fig. S1).

### Correlation of competitive fitness and growth rate

Because it is not technically feasible in this system to measure competitive fitness for the whole experiment, we measured the competitive fitness of a subset of the evolved lineages at one time point, after ~1,000 or ~3,500 generations (for populations evolved at 24°C or 37°C, respectively) and compared this fitness metric to our measurements of growth rate. Competitive fitness was measured in replicate by competing a GFP labeled strain (Cui et al. 2006) against the experimental strain. The two strains were mixed in approximately 1:1 ratios and the density of both strains was determined using a flowcytometer. The culture was allowed to grow overnight at room temperature after which time the flow-cytometer was used again to measure the ratio of the two strains. Competitive fitness was calculated by dividing the natural log of the ratio of the final population density to the initial population density of one strain by the natural log of the ratio of the final population density to the initial population density of the other strain (Wiser and Lenski 2015). Competitive fitness estimates correlated with our growth rate estimates (Pearson’s correlation coefficient = 0.7999; Fig. S2) indicating that growth rate is a good proxy for fitness.

### Data analysis

~36,000 growth curves were collected from all populations over the first 4,000 generations of evolution. This provided us with ~1,500 growth rate estimates per population over this period, approximately half at each temperature.

A generalized additive mixed model (GAMM; see supplementary information section Table S10 for more detail) was fit to the mean growth rate of each population per plate assayed in the environment in which they evolved. Growth rate was fit as a function of generations. Models were fit that included various combinations of the terms genotype, temperature, and generations and the AICs were compared using evidence ratios (ER = e^(0.5*Δ*AIC*)) to assess the significance of terms, including pair-wise and three-way interactions. The three-way interaction relates to the way differences among the genotypes change differently at either temperature. In other words, are there differences in the patterns of convergence or divergence among genotypes between the two temperatures? We also fit a standard least square model (see supplementary material Table S11 for more detail) to the same dataset to assess the effects of each of the parameters used in our GAMM fit.

We fit hyperbolic, power law, and linear models to the growth rate trajectories of all populations assayed in the environment in which it evolved (model details are in Table S4). This analysis was performed on the mean growth rate of each population per plate. We computed the AICc of each fit and calculated the evidence ratio (ER = e^(0.5*Δ*AIC*)) to determine which model (hyperbolic, power law, or linear) best fit the trajectory.

To assess specific time points, as well as for simplicity in visualization, growth rate data were also binned into 250-generation intervals (generation 0 = 0-125, generation 250 = 125-375, generation 500 = 375-625, etc.) and the mean growth rate at both temperatures for each population was calculated. For each population the bin with the highest growth rate for either temperature was identified and the absolute (i.e., maximum mean population growth rate in a 250-generation bin minus the growth rate of the ancestor of that population) and the mean relative increase (i.e., (absolute increase/ancestral growth rate) x 100) in growth rate was calculated from this. ANOVAs testing the effects of genotype, assay temperature, and evolution temperature were performed on these data (*absolute increase/relative increase in growth rate ~ genotype, assay temperature, evolution temperature, genotype*assay temperature, genotype*evolution temperature, assay temperature*evolution temperature*; Tables S2 and S3). For each ANOVA, the residuals were checked for heteroscedasticity both visually and by regression analysis and none was detected. ANOVAs were also performed separately on the 48 data points (24 populations x 2 assay temperatures) in each bin to test for the effect of assay temperature, evolved temperature, genotype, and their interactions as evolution progressed (*mean population growth rate in 250-generation bin ~ genotype, assay temperature, evolution temperature, genotype*assay temperature, genotype*evolution temperature, assay temperature*evolution temperature*; Tables S5, S6, and S7). A Wilcoxon test was also used to test for significant differences between genotypes (Fig. 3)

**Figure 3.**
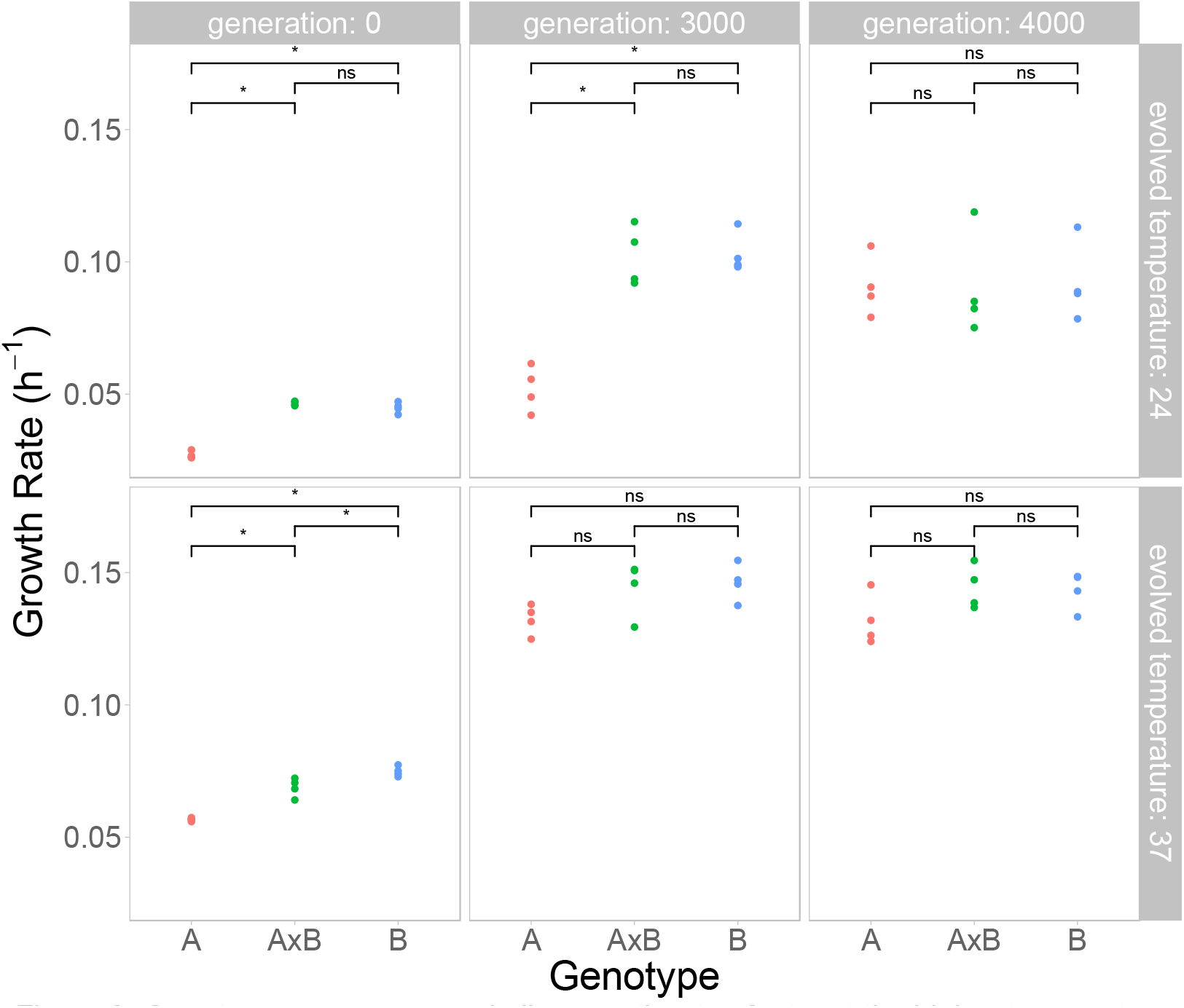
Genotypes converge on similar growth rates faster at the higher temperature. Differences in growth rates in the home environment (i.e. assay temperature the same as the evolution temperature) among genotypes (A = red, AxB = green, B = blue) are shown at three time points (0, 3,000, and 4,000 generations) at each temperature. Each point shows the mean growth rate of one out of the four replicate populations. A Wilcoxon test was used to determine significant differences between genotypes (“*” indicates p < 0.05, “ns” indicates no significant difference).

To test for significant differences at specific time points among populations evolved from a single ancestor nested ANOVAs were performed on the binned data. This analysis (*mean growth rate/plate ~ genotype, replicate population[genotype]&Random, assay temperature, genotype*assay temperature*; Table S8 and S9) tested the effects of replicate population treated as a random effect and nested within genotype, genotype, assay temperature, and the interaction between genotype and assay temperature on the mean growth rate of each population per plate. To test for differences in the variance among replicate populations between evolution temperatures, ANOVAs were performed separately for each evolution temperature. This analysis (*mean population growth rate/plate ~ genotype, replicate population[genotype]&Random*; Fig. 4) tested for effects of replicate population treated as a random effect and nested within genotype and genotype on the mean growth rate of each population per plate in the evolution environment. From this, variance components attributable to replicate population were computed to assess the amount of variation that results from differences among replicate populations; the inverse of this was our measure of repeatability. The same analysis was performed without nesting replicate population in genotype to assess the total variance among all populations as evolution progressed (*mean population growth rate/plate ~ population&Random*; Fig. 5). This analysis shows how the variation between replicates within a genotype interacts with the variation that results from differences between genotypes. At each binned time point, Levene’s tests were performed to assess the significance of differences between evolution temperatures in the variation in growth rate generated by differences both among replicate populations of a single starting genotype and among all populations regardless of genotype.

**Figure 4.**
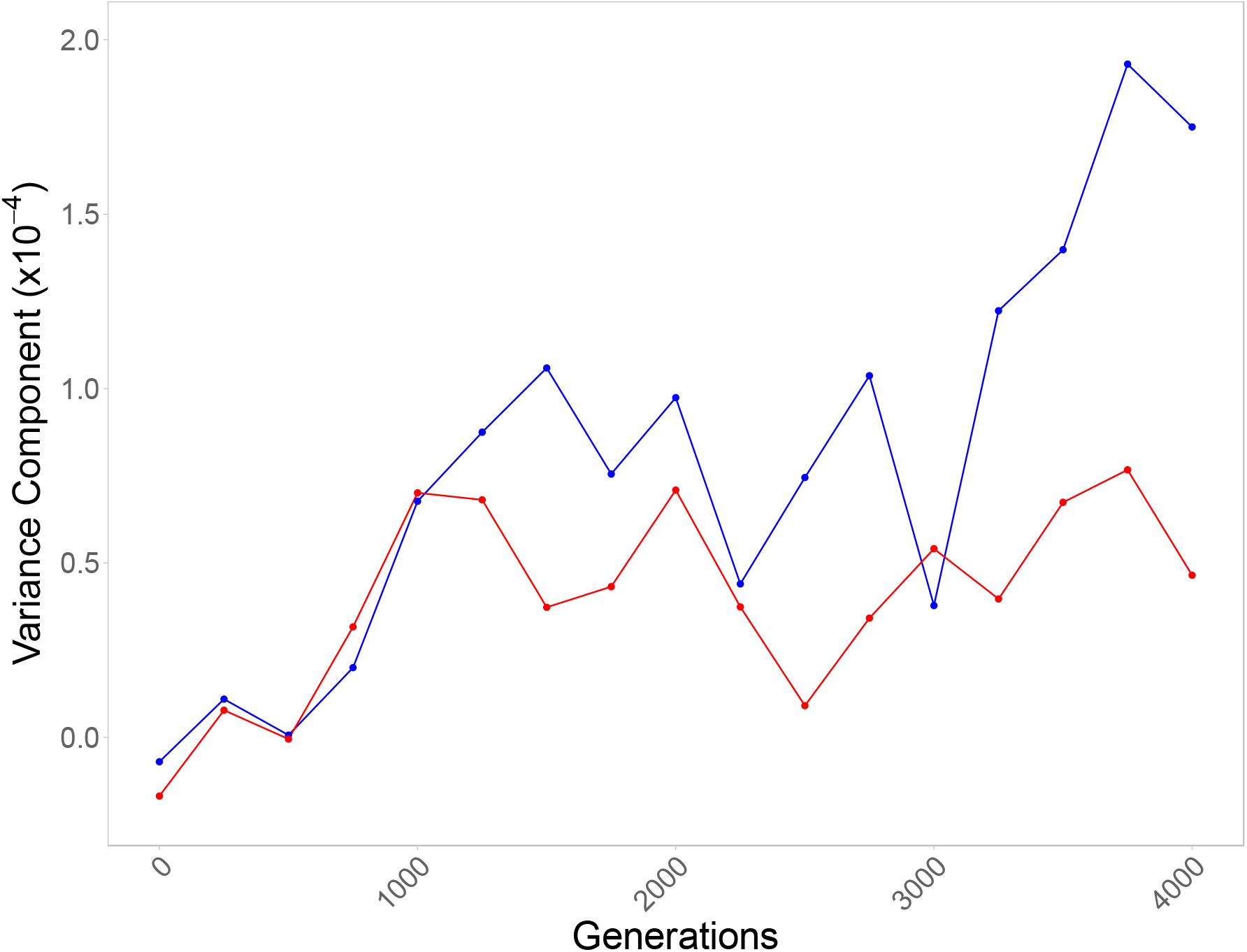
Variance in growth rate due to divergence among replicate populations. The variance components attributable to replicate population for populations evolved and assayed at 24°C (blue) or 37°C (red) over 4,000 generations of evolution. Variance components were estimated from an ANOVA with replicate population nested within genotype (*mean population growth rate/plate ~ genotype, replicate population[genotype]&Random*) for each 250-generation bin and evolution temperature.

**Figure 5.**
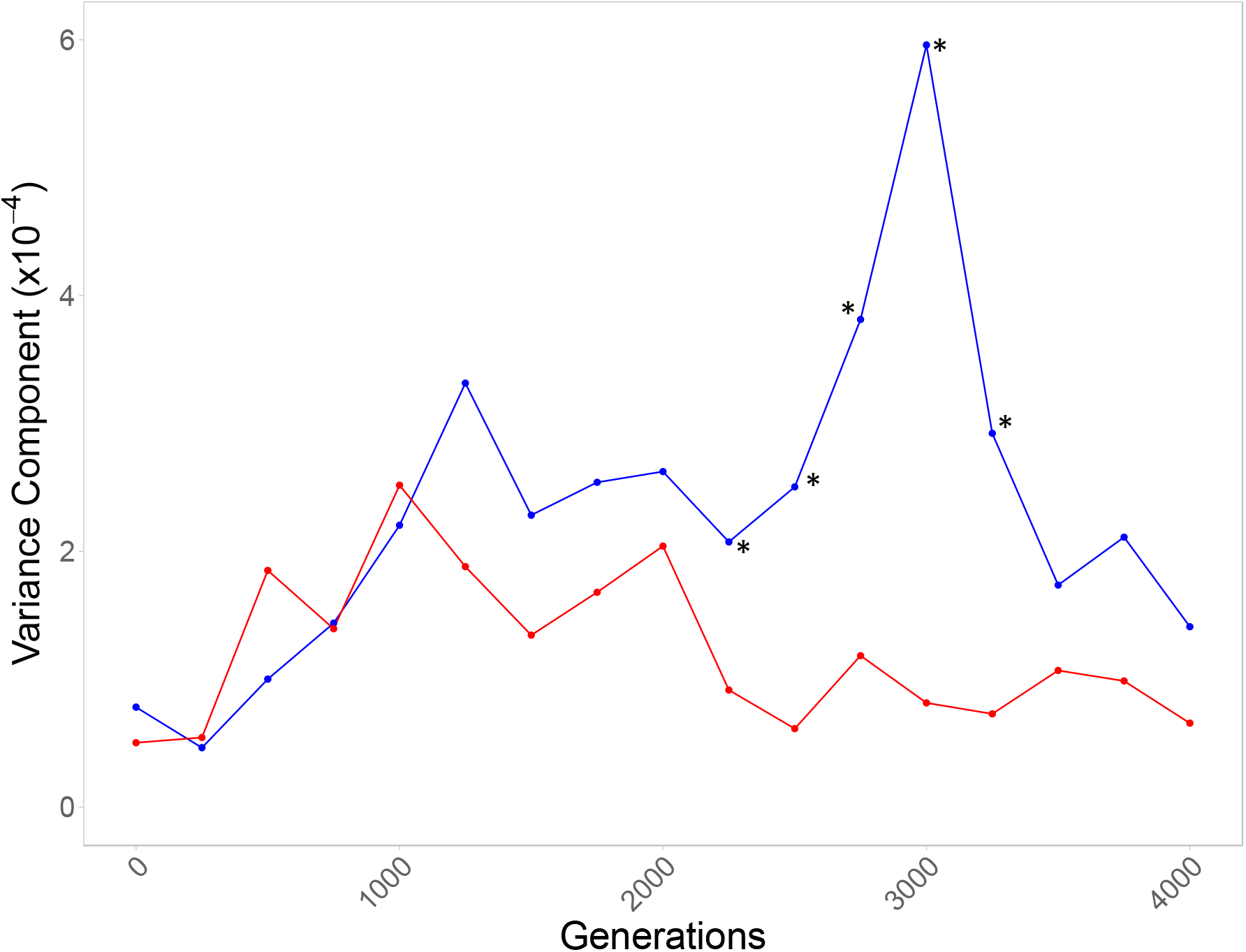
Variance in growth rate among all populations is lower for the hotter populations. The variance components attributable to population for populations evolved and assayed at 24 °C (blue) or 37 °C (red) over 4000 generations of evolution. Variance components were estimated from an ANOVA without population nested within genotype (*mean population growth rate/plate ~ population&Random*) for each 250-generation bin and evolution temperature. Asterisks indicate significant results of Levene’s test.

## Results

### General patterns of adaptation

All populations showed the expected pattern of increased growth rate over the course of the experiment. The trajectories of evolving laboratory populations often follow a pattern of a decelerating rate of return, characterized by larger fitness increases early in the experiment, followed by incrementally smaller increases in subsequent generations (Couce and Tenaillon 2015; Schoustra et al. 2016; Wünsche et al. 2017). Our results follow this pattern (with a linear model fitting the trajectories poorly; Table S4) and this appears consistent at both temperatures (Fig. 1) and in all three genotypes when analyzed separately (Fig. 2), suggesting that experimental evolution in the ciliate *T. thermophila* does not fundamentally differ from other taxa.

**Figure 1.**
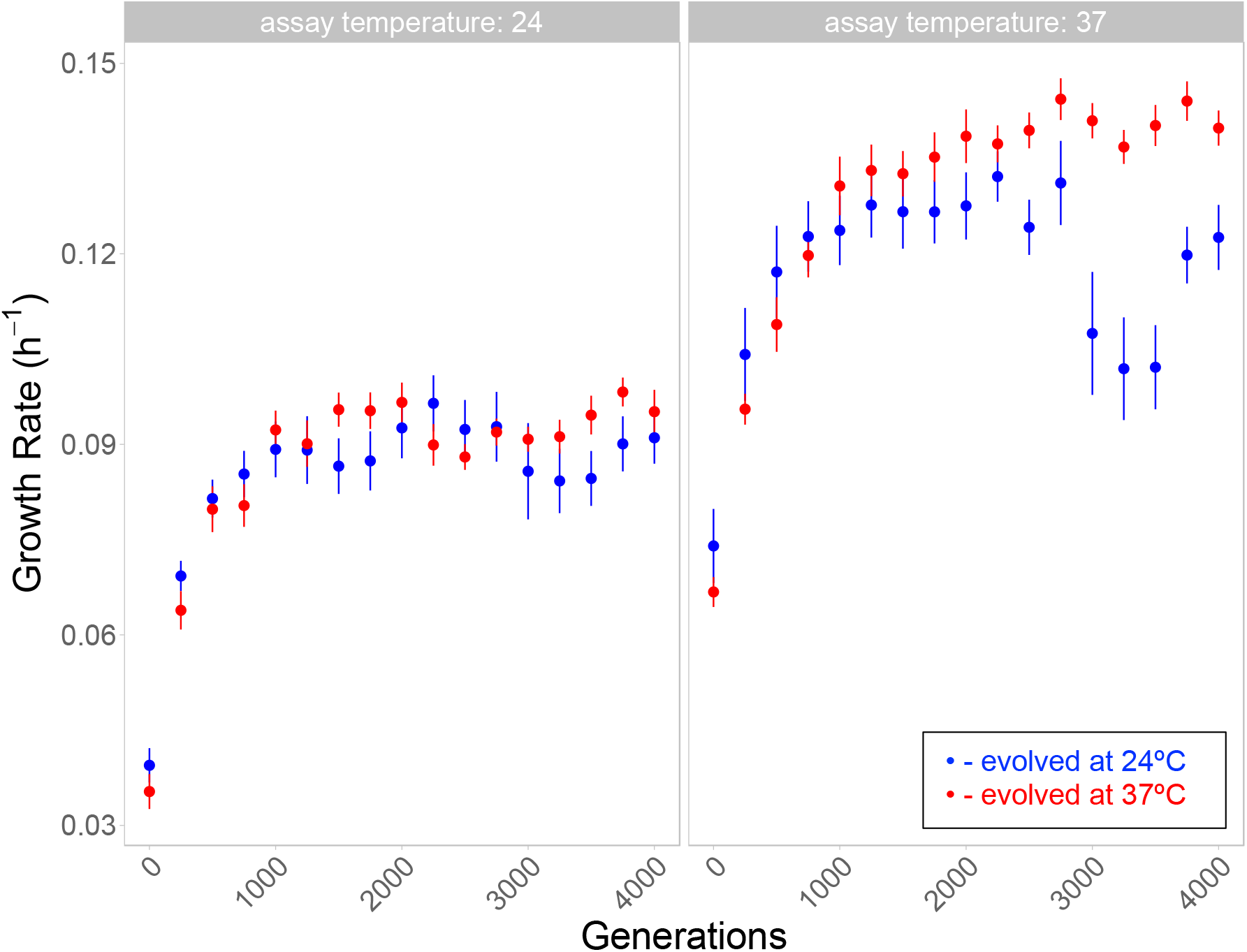
Overall pattern of evolution across all populations assayed at 24°C and 37°C. Mean growth rate and 95% confidence intervals of populations evolved at 24°C (blue) and 37°C (red) when assayed at 24°C (left panel) and 37°C (right panel) are shown over 4,000 generations. Data are binned into 250 generation intervals, with the first bin containing generations 0-125.

**Figure 2.**
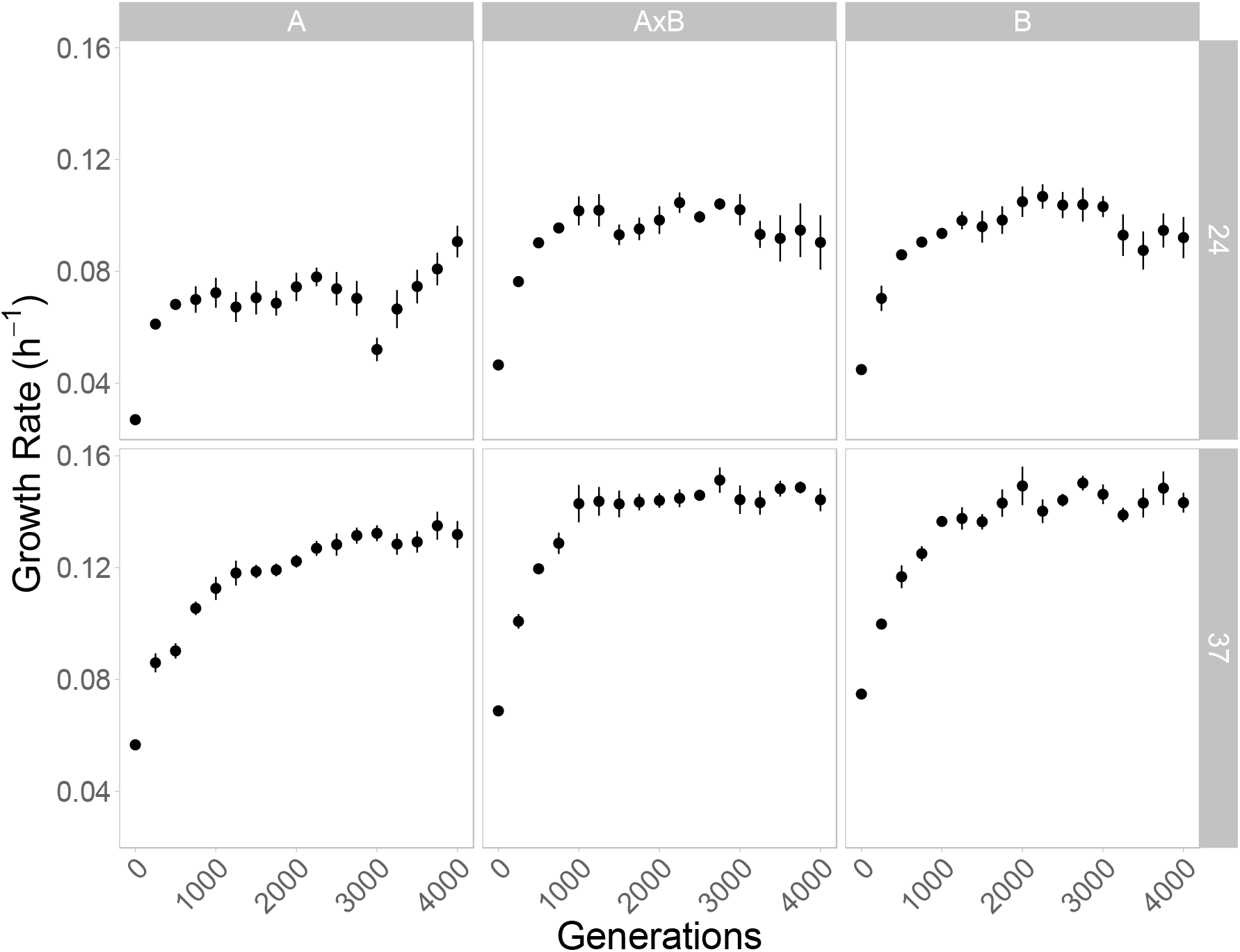
Fitness trajectories of each genotype assayed in their evolved temperature (correlated response at alternative temperature not shown). Mean growth rate and 95% confidence intervals of four replicate populations for each genotype are shown over 4,000 generations. The top panels show populations evolved and assayed at 24°C and the bottom shows populations evolved and assayed at 37°C. Data are binned as in Fig. 1.

Previous experiments have also shown that populations founded by initially slower growing genotypes tend to increase more in growth rate over the course of an experiment than those founded by initially faster growing genotypes (Jerison et al. 2017; Wünsche et al. 2017). We found a qualitatively similar result whereby genotype had a significant effect on the absolute increase (ANOVA: *F* (2,38) = 4.48, *P* = 0.0179; Table S2) and the relative increase (ANOVA: *F* (2,38) = 192.39, *P* < 0.0001; Table S3) in growth rate, and with populations founded by the slowest growing genotype (A) experiencing the largest increases in growth rate for all four combinations of evolution temperature and assay temperature. The mean absolute increase (i.e., the mean growth rate from the highest recorded 250-generation bin minus the growth rate of the ancestor of that population) and the mean relative increase (i.e., (absolute increase/ancestral growth rate) x 100) in growth rate are reported for each combination of genotype, evolution temperature, and assay temperature in Tables S1a and S1b. We also calculated the scaled effect of parent based on the best fit model identified by GAMM (see below) and found that parent A by generations was significantly positive, while the scaled effect of parent B by generations and AxB by generations was negative, supporting the hypothesis that the slowest growing genotype experiences the greatest increase in growth rate (Table S11). However, due to the small number of genotypes (3) used in this experiment we cannot definitively say this effect is due to the initially lower starting growth rate of genotype A.

Unlike the long-term evolved *E. coli* lines, which continue to increase in fitness even after 60,000 generations (Lenski et al. 2015), we find no significant change in mean growth rate among populations over the most recent 1,000 generations of evolution; in fact our estimate of mean growth rate drops slightly from 0.1151 divisions per hour (h^-1^) at 2750 generations to 0.1130 h^-1^ at 4,000 generations. Additionally, a hyperbolic model yields a substantially better fit than a power law model or a linear model, generating a significantly lower AIC value (Table S4). This suggests that the populations may have reached growth rate optima upon which further improvement is unlikely. However, given the limited number of generations and smaller population sizes, we are cautious in interpreting this result as further evolution could lead to increases in growth rate altering our model fits. It is also important to consider that fitness could be increasing in ways that are not captured by our growth rate estimates so that growth rate may have plateaued while fitness is still being optimized in other ways e.g., increase in carryingcapacity or decrease in lag-time (Li et al. 2018).

### Evolution at a higher temperature results in faster convergence among genotypes

At the start of the experiment there was a significant difference in growth rate between genotypes (ANOVA: *F* (2,38) = 189.38 *P* < 0.0001; Table S5). This was true whether populations were assayed at 37°C or 24°C (Wilcoxon tests; Fig. 3). Specifically, one of the parental genotypes (A) grew significantly slower than the other parental genotype (B) and the hybrid genotype (A×B) at both temperatures.

To determine which factors affect the evolutionary trajectories of the different populations of these genotypes, we fit a GAMM and found that including the three-way interaction between genotype, temperature, and generation produced the best fit with the lowest AICc (see Table S10 for the models fit, the AICc of each model, and the evidence ratios indicating the superior fit of the model that included the three-way interaction). Based on this result, we fit a standard least square model using the same terms (generations, genotype, temperature, and all interaction terms) and found that the scaled effect of generations by slower growing parent (A) by 24°C was significantly negative while the effect of generations by slower growing parent (A) by 37°C was significantly positive (Table S11). This result indicates that genotypes are converging faster at the higher temperature.

To further explore this result, we used ANOVA to determine at which generations there remains a significant difference between genotypes at each temperature. The difference between genotypes remained at both temperatures for nearly 3,000 generations of evolution. After 3,000 generations, we still find an effect of genotype on growth rate (ANOVA: *F* (2,38) = 14.79, *P* < 0.0001; Table S6), however after investigating the significant interaction effect of genotype by evolution environment (ANOVA: *F* (2,38) = 6.21, *P* = 0.0047; Table S6) we found this effect is driven primarily by the 24°C evolved populations at this time point. In fact, the significant difference between genotypes is lost after 3000 generations of evolution at 37°C (R^2^ = 0.0301) but not at 24°C (R^2^ = 0.472; Wilcoxon test; Fig. 3), supporting the finding that the genotypes converge on a similar growth rate more quickly at the higher temperature. By 4,000 generations there is still a significant, but smaller effect of genotype on growth rate (ANOVA: *F* (2,38) = 3.44, *P* = 0.0425; Table S7) however Wilcoxon tests detect no significant differences between genotypes at either temperature (Fig. 3).

### Evolution at a higher temperature results in less variation among replicate populations

The variation in growth rate among replicate populations appeared greater in populations evolved at 24°C compared to those evolved at 37°C. To test whether apparent differences between replicate populations evolved from a single ancestor were significant we performed a nested ANOVA on mean growth rate per plate at 4000 generations. We found a significant effect of replicate population nested within genotype (F (21,826) = 13.95, P < 0.0001; Table S8) indicating significant divergence between populations evolved from a single ancestor. Similar results were obtained for other time points. In fact, even as soon as generation 125 there is an effect of population nested within genotype (F (21,283) = 2.65, P = 0.0002; Table S9) indicating that populations began to evolve measurable differences in growth rate early in their evolution. To further analyze this result and to assess differences in the variance produced at either evolution temperature, we performed Levene’s test every 250 generations and compared the variance component attributable to replicate population (nested within genotype) at either evolution temperature (Fig. 4). The variance component attributable to population is a measure of repeatability because it describes how similar or different the growth rates of replicate populations are within each genotype. We also compared the variance component attributable to population regardless of genotype using an unnested model for either temperature (Fig. 5). This allows us to see how the decrease in variation between genotypes (Fig. 3) interacts with the variation produced among replicate populations of a given genotype (Fig. 4) to affect the overall variation between all populations regardless of genotype.

The small sample size within a genotype (n=4) meant Levene’s test was unable to detect significant differences in the variance between temperatures at each individual time point, but we consistently see a larger variance component attributable to replicate population nested within genotype among populations evolved and assayed at 24 °C particularly after 1000 generations (Fig. 4). This is true regardless of assay temperature, indicating that evolution temperature is likely driving this effect, and supporting our hypothesis that temperature impacts the repeatability of the growth rate trajectories of replicate populations.

When we combine growth rate data from all genotypes Levene’s tests indicate there is a significant difference in the variance among populations at either temperature from generation 2,250 to generation 3,250 (Fig. 5). We also find consistently lower variance components attributable to population among 37°C-evolved populations than those evolved at 24°C (Fig. 5). This is due to the joint effect of less divergence between replicate populations of the same genotype (Fig. 4) and more convergence among different genotypes for populations evolved at 37°C relative to those evolved at 24°C (Fig. 3). At both temperatures the variance component attributable to population peaks at an intermediate generation, although the peak is higher and later for populations evolved at 24°C, as variation accumulates among replicate populations but before genotypes have had sufficient time to converge (Fig. 5).

In spite of the greater variation among replicate populations of the same genotype evolved at 24°C (Fig. 4) we still detect greater differences among genotypes when evolution takes place at 24°C (Fig. 3). This indicates that the observed differences among genotypes at 24°C vs. 37°C (described in the section above) are not just due to higher variability among replicate populations at the lower temperatures, but also to longer lasting differences between genotypes. Additionally, the increased variance among lines evolved at the colder temperature is consistent when we look at the growth rate at the alternate temperature indicating this pattern is not the result of measurement differences between the two temperatures and is indeed the result of the evolution temperature.

### Asymmetry of the correlated responses

By generation 4,000, all populations increased in growth rate at both the temperature in which they evolved and the alternate temperature (Table S1), indicating no evidence of trade-offs at this time point. However, we find a marginally significant interaction between evolution temperature and assay temperature (ANOVA: *F* (1,38) =3.17, *P* = 0.0829; Table S7) at generation 4,000. This suggests that some of the adaptation that has taken place over the course of the experiment is temperature-specific despite an overall correlation between growth rates of evolved populations at either temperature (r = 0.597). This correlation is even greater when the ancestors are included in the analysis (r = 0.858; Fig. 6).

**Figure 6.**
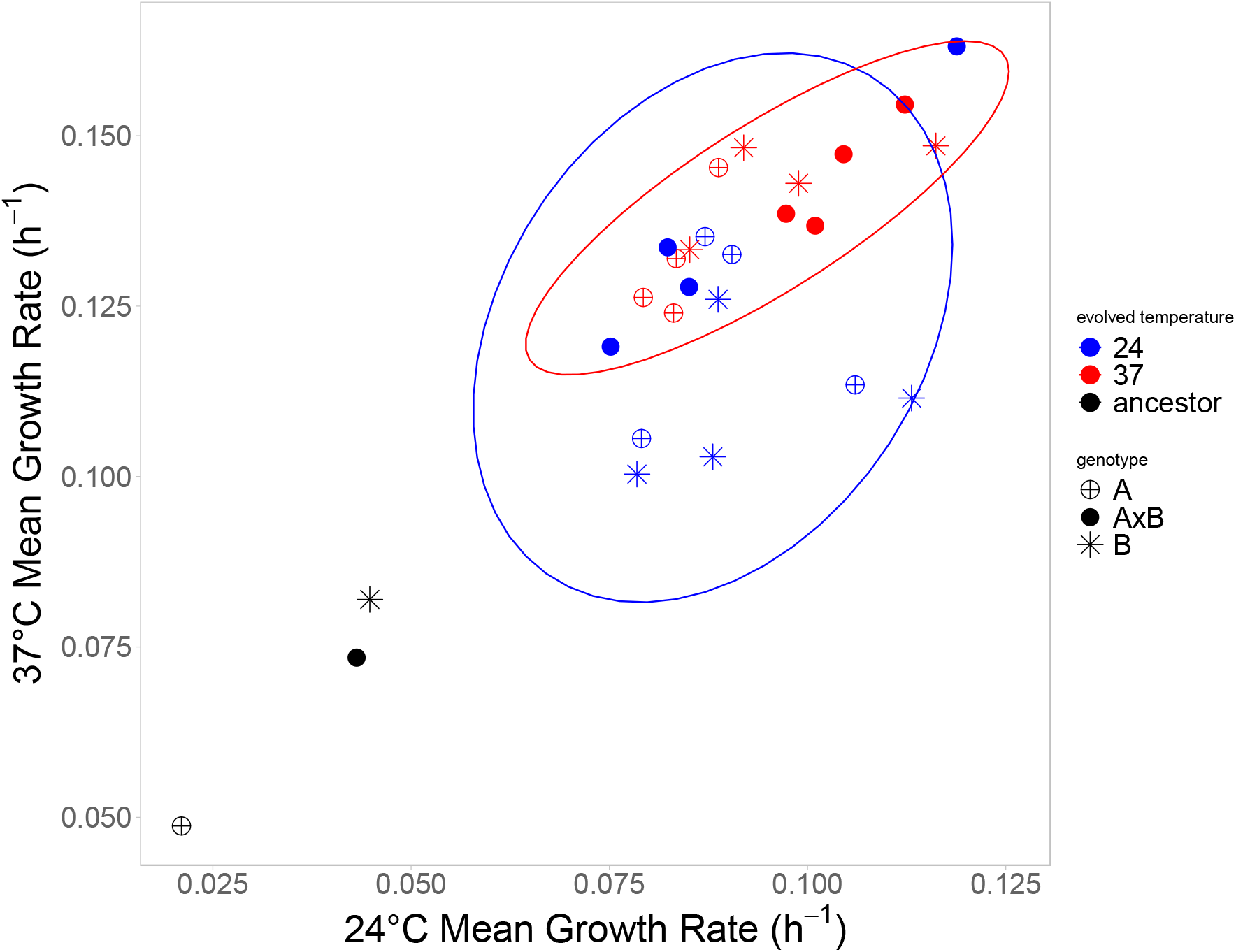
Correlation between growth rates in alternative environments. Growth rate of populations after 4000 generations of evolution, measured at 37°C (y-axis) or 24°C (x-axis). Genotypes are indicated by the symbols and the evolution environment is indicated by red (37 °C) or blue (24 °C) with the ancestors shown in black. A trade-off exists if an evolved population has lower fitness than its ancestor at the alternate temperature from which it evolved. No tradeoffs are observed here. The 95% confidence ellipse is shown for populations evolved at 37 °C (red) and for populations evolved at 24 °C (blue).

To assess which temperatures were driving the interaction between evolution temperature and assay temperature, we compared growth rates from each assay temperature. We found a significant effect of evolution temperature when assays were performed at 37°C (R^2^ = 0.285) but, remarkably, not at 24 °C (R^2^ = 0.0265; Tukey-Kramer: p < 0.05). This means that even after 4,000 generations of evolution, the temperature at which populations evolved makes no difference when growth rate is assayed at 24 °C. This indicates there is a greater correlated response when evolution occurs at 37 °C. In other words, evolution at the hotter temperature increased growth at the colder temperature more than evolution at the colder temperature increased growth at the hotter temperature (Fig. 6).

## Discussion

We examined the evolutionary trajectories of populations of different genotypes of *T. thermophila* under differing temperature regimes. Our experimental design allowed us to test how evolution temperature affects repeatability, as well as how it impacts historical differences as evolution progressed at each temperature. We found that the hotter temperature resulted in greater repeatability of evolution and faster convergence between divergent genotypes.

After 4,000 generations, we found that populations evolved at 37 °C significantly outperformed those evolved at 24 °C (Fig. 1). This outcome aligns with previous findings that “hotter is better” (Knies et al. 2009; Angilletta et al. 2010). This hypothesis states that hot-adapted genotypes will have higher maximum growth rates than cold-adapted genotypes because they have evolved greater robustness in response to the chemical and metabolic reactions happening more quickly at hotter temperatures and because the rate-depressing effects of low temperature cannot be overcome by adaptation or plasticity.

### Temperature affects the convergence of different genotypes

Over the course of evolution, different starting genotypes and phenotypes could converge, evolve in parallel, or diverge even further. Through epistatic interactions, genotype can constrain the future evolution of a population by biasing the set of available beneficial mutations that are likely to be selected (Draghi and Plotkin 2013). Similar genotypes are expected to fix a similar set of mutations while more divergent genotypes are expected to fix a less similar set of mutations leading to further divergence between the genotypes (Blount et al. 2018; Starr et al. 2018). At the same time natural selection could overcome both random drift and epistatic interactions to produce convergence between divergent genotypes.

Previous experiments have found that the rate of adaptation is inversely proportional to initial fitness and that initially different populations often end up at the same fitness optima (Jerison et al. 2017; Wünsche et al. 2017). At the same time studies have also found that particular alleles can impede this fitness recovery and constrain the future of evolution (Woods et al. 2011; Jerison et al. 2017). However, these experiments were limited to less than 1,000 generations of evolution and it is unclear whether continued evolution would eventually allow these populations to reach the same fitness optimum as their relatives. For more distantly related populations, we might expect this process to take longer if it even occurs at all.

In our experiment, the maintenance of historical differences between divergent genotypes of the same species over many generations of evolution at both temperatures suggests that genetic differences in the initially slowest growing genotype are impeding future adaptation in a manner that is not easily overcome. Despite the overall increase in growth rate being greatest for the initially less fit genotype as expected, we observe slower rates of adaptation for this genotype than we would expect if all genotypes followed the same pattern of diminishing returns epistasis. We also find that temperature affects this pattern and the rate of convergence. Differences in growth rate between genotypes were maintained for over 3,000 generations at 24°C while convergence among the genotypes was more rapid at 37°C. Why a higher temperature would be more conducive to convergence is unclear but could be related to other effects of temperature observed in our experiment. For example, higher selection coefficients and/or more targets of selection at 37°C may contribute to the slower growing genotype catching up more quickly at this temperature, to the greater repeatability, and to the asymmetry of the correlated responses.

The ability of populations to escape constraints on evolutionary change can be vital to long-term survival (Chao and Weinreich 2005; Weinreich et al. 2005). In this experiment, we show the gradual loss of growth rate differences between genotypes even while differences evolve among replicate populations of the same genotype at both temperatures. This suggests that differences in patterns of divergence depend on relatedness, e.g. increasing divergence among genetically identical replicates, but decreasing variation among less related genotypes as the mean growth rates of divergent genotypes converge in the same environment. However, very distantly related genotypes may find drastically different evolutionary solutions to the same environmental pressures, which could contribute to further phenotypic divergence. Therefore, it is possible that phenotypic divergence is minimized at intermediate levels of relatedness.

### Temperature affects repeatability among populations

Previous studies have found differences in the repeatability of evolutionary trajectories under different environmental conditions (e.g., Gresham et al. 2008; Bailey et al. 2015). In these experiments, replicate populations were more likely to diverge in some environments but experience repeatable evolutionary trajectories in others. Likewise, we found that replicate populations of all genotypes diverged more at 24°C and were more repeatable at 37°C.

The greater variation among populations evolved at 24°C suggests that these evolutionary trajectories are more dependent on chance events than the populations evolved at 37°C. This result may reflect differences in the environment that affect the degree of epistasis or “ruggedness” of the fitness landscape and/or rate of mutation and distribution of their effects.

Differences in the “ruggedness” of the fitness landscape, caused by epistatic interactions (Kvitek and Sherlock 2011; Poelwijk et al. 2011), at each temperature could explain our observation of increased repeatability at 37°C. While theory predicts that a rugged fitness landscape can increase the repeatability of evolution at the level of the mutational pathways followed (De Visser and Krug 2014) the opposite is true at the fitness level (Bank et al. 2016). Therefore, theory suggests, the greater repeatability in growth rate (a good proxy for fitness) trajectories at 37°C could result from a more uniform fitness landscape at this temperature.

Greater repeatability could also result from a difference in the distribution of beneficial mutations available in each environment (Lenski et al. 1991). At 24°C, the lower repeatability suggests there may be rare highly beneficial mutations that increase growth rate in some but not all populations, while at 37°C there may be fewer of these types of mutations resulting in growth increasing more uniformly across replicate populations. If this were the case, we would eventually expect to see a reduction in the variation among replicate populations evolved at 24°C. Continued experimental evolution of our populations may eventually lead to this result, but if epistatic interactions are important, as they appear to be (Kuzmin et al. 2018), they may constrain future evolution making eventual convergence even more unlikely.

The strength of selection may also differ in these environments. Theoretical results suggest that stronger selection results in increased repeatability (Orr 2005). This theory is corroborated by a meta-analysis showing a strong positive relationship between population size, with larger populations experiencing greater selection, and greater repeatability (Bailey et al. 2017). Our populations are approximately the same size at either temperature meaning our observations are not simply a reflection of differences in the sizes of the populations at either temperature. However, 37°C is near the upper limit of the thermal tolerance for this species (Hallberg et al. 1985), which may pose a greater selective pressure thereby causing the observed reduction in variation among populations evolved at this temperature.

### Temperature affects correlated responses

Experiments using *E. coli* have found substantial evidence for temperature associated trade-offs (Bennett et al. 1992; Bennett and Lenski 1993, 2007; Mongold et al. 1996; Woods et al. 2006). In *T. thermophila*, we find no evidence for trade-offs in any of our populations after 4000 generations. However, we do find an asymmetric correlated response, whereby evolution at 37°C increases growth rate at 24°C more than evolution at 24°C increases growth rate at 37°C, which is similar to what is observed in *E. coli*. Evolution at a hotter temperature increases growth rate at a colder temperature for both species while evolution at a colder temperature increases growth rate at a hotter temperature less for *T. thermophila* and often decreases it for *E. coli* (Bennett et al. 1992; Bennett and Lenski 1993; Mongold et al. 1996). One likely explanation for the difference between *T. thermophila* and *E. coli* is that the *E. coli* experiments started from an ancestor that had already evolved under laboratory conditions for 2,000 generations and was therefore pre-adapted to the general culture conditions, as opposed to our *T. thermophila* lines, which were derived from wild collected strains grown in lab only ~500 generations before cryopreservation. Thus, it seems likely that a greater proportion of the adaptation that occurred in the *T. thermophila* populations, compared to the *E. coli* populations, involved adaptation to the general culture conditions as opposed to the specific temperature.

As evolution occurs in one environment, fitness may change in other environments either as a direct pleiotropic response to selection in the evolution environment or due to the accumulation of mutations that are neutral in the evolution environment but have fitness consequences in the other environment (Cooper and Lenski 2000). The asymmetry we observe in the correlated responses could be due to asymmetry in the pleiotropic responses, whereby a 37°C beneficial mutation increases growth rate more at 24°C than a 24°C beneficial mutation does at 37°C. Alternatively, the asymmetry in the correlated responses could arise from an asymmetry in the effect of neutral and nearly neutral mutations at the alternate temperature. In other words, the neutral and nearly neutral mutations that are able to accumulate at 37°C are also mostly neutral at 24°C while the neutral and nearly neutral mutations that are able to accumulate at 24°C tend, on average, to be slightly deleterious at 37°C. These two possibilities are not mutually exclusive.

One possible mechanistic explanation for the observed asymmetry could be more transcript diversity, and thus more targets of selection, in hotter conditions resulting in most genes that are transcribed and selected at 24°C also being transcribed and selected at 37°C but not vice versa. This would be consistent with the lack of antagonistic pleiotropy across temperatures among the most positively selected mutations found in lab-evolved *E. coli* (Deatherage et al. 2017) and is supported by data showing that more genes are up-regulated at hotter temperatures (Tai et al. 2007; Mittal et al. 2009). Additionally, the 37°C evolved populations divide more quickly and experience a greater density range, and thus a more heterogenous environment, than those evolved at 24°C, which could also contribute to greater transcript diversity and the asymmetry in the correlated response that we observe. This idea is supported by a meta-analysis of trade-off experiments, which found that populations evolved in homogeneous environments exhibited more trade-offs than populations evolved in temporally heterogeneous environments (Bono et al. 2017). However theoretical predictions made by Gilchrist (1995) suggest, somewhat counterintuitively, that the opposite should be true and that temporal heterogeneity should lead to greater thermal specialization. The 37°C populations also experience an additional possible source of heterogeneity because the 37°C tubes are not pre-heated so the cells experience the 24°C temperatures for a very brief period each day. It is conceivable that this very brief period of cold is sufficient to explain the greater correlated response in the 37 °C evolved populations. However, we consider this unlikely as this cold exposure is taking place during lag phase, not when cells are dividing, and is therefore unlikely to impact selection on the growth rate.

The asymmetric correlated response we observe may also be related to the other effects of evolution temperature that we observed. For example, the conditions responsible for greater convergence and repeatability when evolution occurs at 37°C may also act to optimize and constrain growth rate at the lower temperature. Thus, our results are consistent with there being more targets of selection at 37°C, which would lead to faster adaptation, greater repeatability, and asymmetric correlated responses. It is also possible that all of these results are a reflection of the “hotter is better” hypothesis (Knies et al. 2009; Angilletta et al. 2010). However, this hypothesis does not directly explain the observed correlated responses of evolution in hotter conditions indicating that different aspects of the 37°C environment may be responsible for the greater convergence, the greater repeatability, and the larger correlated response. In the future, more high-throughput methods with greater control of the evolution conditions will allow for the identification of the precise environmental conditions responsible for the difference that we observed in evolution at different temperatures.

Another possible interpretation of our results is that populations evolving at 24°C adapt by increasing different components of fitness than those evolving at 37°C. We measured growth rate, which is a major component of fitness, and well correlated with competitive ability in our experiments, but fitness can also increase in more complex ways than simply increasing maximum growth rate (Li et al. 2018). For example, decreasing lag time or increasing carrying capacity could increase fitness without affecting growth rate. Additionally fitness gains can be accrued and realized in different portions of the growth-cycle (Li et al. 2018), which could contribute to the asymmetry of the correlated responses that we observe if the amount of time spent in different phases of the growth cycle differs substantially between temperatures. A final caveat is that all of the adaptation that we observed occurred in the somatic nucleus, which is discarded following sexual reproduction. While there is evidence of some epigenetic inheritance between parental and progeny somatic genomes (Beisson and Sonneborn 1965; Chalker and Yao 1996; Pilling et al. 2017), it is unknown whether any of the adaptation that occurred in our experimental populations would be inherited by newly produced sexual progeny. However, this may be a moot point in this experiment because all of the evolved populations lost the ability to undergo sexual conjugation, at least under laboratory conditions.

### Conclusion

One of the most important questions for evolutionary biologists is how variation builds up over time to create all of the diversity observed around us. Small incremental changes in isolated populations can, given enough time, lead to major differences in the organisms that make up those populations. However, selection can also result in striking examples of parallel and convergent evolution and we are only beginning to understand the ways in which genotype and the environment contribute to this process and to the overall repeatability of evolution. Here, we demonstrated that the temperature at which populations evolve can affect the patterns of evolution, with populations in hotter environments showing greater repeatability among replicates and faster convergence among genotypes. In addition, evolution at the hotter temperature results in populations that are more fit in the colder temperature than vice versa. These results support the growing body of work that demonstrate the importance of environment in determining evolutionary trajectories of populations.

