## supporting information for "Temperature affects the repeatability of evolution in the microbial eukaryote *Tetrahymena thermophila*"

**Tables**

| | **Evolved at 24°C** | | **Evolved at 37°C** | |
|---|---|---|---|---|
| | Assayed at 24°C | Assayed at 37°C | Assayed at 24°C | Assayed at 37°C |
| Genotype A | **0.064** (0.057-0.071) | **0.078** (0.073-0.084) | **0.072** (0.067-0.077) | **0.087** (0.077-0.097) |
| Genotype B | **0.066** (0.053-0.079) | **0.065** (0.032-0.098) | **0.061** (0.045-0.078) | **0.072** (0.060-0.085) |
| Genotype AxB | **0.067** (0.053-0.082) | **0.072** (0.057-0.087) | **0.063** (0.057-0.068) | **0.080** (0.071-0.089) |
| Overall | **0.066** (0.061-0.070) | **0.072** (0.063-0.080) | **0.065** (0.060-0.070) | **0.080** (0.074-0.086) |

Table S1a. Mean increase in absolute growth rate ($H^{-1}$) for each genotype, evolution environment, and assay temperature with 95% confidence intervals. The mean increase of all 12 populations regardless of genotype is shown in the final row of the table.

| | **Evolved at 24°C** | | **Evolved at 37°C** | |
|---|---|---|---|---|
| | Assayed at 24°C | Assayed at 37°C | Assayed at 24°C | Assayed at 37°C |
| Genotype A | **254%** (226-283%) | **151%** (140-163%) | **287%** (267-308%) | **169%** (150-188%) |
| Genotype B | **148%** (119-178%) | **78.9** (39.0-119%) | **138%** (101-175%) | **87.8%** (72.5-103%) |
| Genotype AxB | **157%** (124-191%) | **93.8%** (74.4-113%) | **146** (134-159%) | **104%** (91.9-116%) |
| Overall | **187%** (153-220%) | **108%** (85.2-131%) | **191%** (144-237%) | **120%** (96.2-144%) |

Table S1b. Mean relative increase in growth rate ($H^{-1}$) for each genotype, evolution environment, and assay temperature with 95% confidence intervals. The mean relative increase of all 12 populations regardless of genotype is shown in the final row of the table.

| Source | DF | Sum of Squares | Mean Square | F Ratio | Prob > F |
|---|---|---|---|---|---|
| **Model** | 9 | 0.00273 | 0.000303 | 3.8781 | 0.0015 |
| genotype | 2 | 0.000700 | | 4.479 | 0.0179 |
| evolved temperature | 1 | 0.000177 | | 2.262 | 0.1408 |
| evolved temperature*genotype | 2 | 0.000138 | | 0.8801 | 0.423 |
| assay temperature | 1 | 0.00131 | | 16.69 | 0.0002 |
| genotype*assay temperature | 2 | 0.000201 | | 1.285 | 0.2883 |
| evolved temperature*assay temperature | 1 | 0.000209 | | 2.667 | 0.1107 |
| **Error** | 38 | 0.00297 | 0.000078 | | |
| **Corrected Total** | 47 | 0.00570 | | | |

Table S2. Factors affecting absolute increase in growth rate. Standard least squares fit of the absolute increase in growth rate of each population over 4000 generations with all factors treated as fixed effects.

| Source | DF | Sum of Squares | Mean Square | F Ratio | Prob > F |
|---|---|---|---|---|---|
| **Model** | 9 | 177206.82 | 19689.6 | 75.8655 | <0.0001 |
| genotype | 2 | 99864.22 | | 192.39 | <0.0001 |
| evolved temperature | 1 | 768.83 | | 2.96 | 0.0934 |
| evolved temperature*genotype | 2 | 1779.34 | | 3.43 | 0.0428 |
| assay temperature | 1 | 66660.82 | | 256.85 | <0.0001 |
| genotype*assay temperature | 2 | 7933.99 | | 15.28 | <0.0001 |
| evolved temperature*assay temperature | 1 | 199.62 | | 0.7692 | 0.386 |
| **Error** | 38 | 9862.28 | 259.5 | | |
| **Corrected Total** | 47 | 187069.1 | | | |

Table S3. Factors affecting relative increase in growth rate. Standard least squares fit of the relative increase in growth rate of each population over 4000 generations with all factors treated as fixed effects.

| Model | AICc | AIC evidence ratio |
|---|---|---|
| linear | -21874.29 | >2.0E+60 |
| power law | -23583.44 | 1.032E+13 |
| hyperbolic | -23643.37 | n.a. |

Table S4. Comparison of power law (*mean growth rate per plate ~ mean growth rate of ancestors (0.0507) +* $\theta_2$ *x generations^* $\theta_1$), hyperbolic (*mean growth rate per plate ~ mean growth rate of ancestor + (*$\theta_1$ *x generations)/(*$\theta_2$ *+ generations)*), and linear (*mean growth rate per plate ~ mean growth rate of ancestor +* $\theta_1$ *x generations*) model fits to the observed changes in growth rate over 4000 generations of evolution. The AIC evidence ratio (ER = e^(0.5*$\Delta AIC$)) indicates the hyperbolic model fits the data substantially better than the power law and linear model.

| Source | DF | Sum of Squares | Mean Square | F Ratio | Prob > F |
|---|---|---|---|---|---|
| **Model** | 9 | 0.0201 | 0.00223 | 144.69 | <0.0001 |
| assay temperature | 1 | 0.0131 | | 847.31 | <0.0001 |
| evolved temperature | 1 | 0.000389 | | 25.22 | <0.0001 |
| genotype | 2 | 0.00583 | | 189.38 | <0.0001 |
| evolved temperature*assay temperature | 1 | 0.0000291 | | 1.890 | 0.1773 |
| genotype*assay temperature | 2 | 0.000264 | | 8.576 | 0.0008 |
| evolved temperature*genotype | 2 | 0.000491 | | 15.92 | <0.0001 |
| **Error** | 38 | 0.0005854 | 0.000015 | | |
| **Corrected Total** | 47 | 0.0206 | | | |

Table S5. Factors affecting mean population growth rate at generation 0. Standard least squares fit of mean growth rate of each population at generations 0-125 with all factors treated as fixed effects.

| Source | DF | Sum of Squares | Mean Square | F Ratio | Prob > F |
|---|---|---|---|---|---|
| **Model** | 9 | 0.0337 | 0.00374 | 13.96 | <0.0001 |
| assay temperature | 1 | 0.0155 | | 57.81 | <0.0001 |
| evolved temperature | 1 | 0.00445 | | 16.61 | 0.0002 |
| genotype | 2 | 0.00793 | | 14.79 | <0.0001 |
| evolved temperature*assay temperature | 1 | 0.00243 | | 9.056 | 0.0046 |
| genotype*assay temperature | 2 | 0.0000353 | | 0.0659 | 0.936 |
| evolved temperature*genotype | 2 | 0.00333 | | 6.21 | 0.0047 |
| **Error** | 38 | 0.0102 | 0.000268 | | |
| **Corrected Total** | 47 | 0.0439 | | | |

Table S6. Factors affecting mean population growth rate at generation 3000. Standard least squares fit of mean growth rate of each population at generations 2875-3125 with all factors treated as fixed effects.

| Source | DF | Sum of Squares | Mean Square | F Ratio | Prob > F |
|---|---|---|---|---|---|
| **Model** | 9 | 0.0213 | 0.00237 | 14.61 | <0.0001 |
| assay temperature | 1 | 0.0174 | | 107.36 | <0.0001 |
| evolved temperature | 1 | 0.00137 | | 8.42 | 0.0061 |
| genotype | 2 | 0.00112 | | 3.44 | 0.0425 |
| evolved temperature*assay temperature | 1 | 0.000515 | | 3.17 | 0.0829 |
| genotype*assay temperature | 2 | 0.000273 | | 0.841 | 0.439 |
| evolved temperature*genotype | 2 | 0.000642 | | 1.98 | 0.152 |
| **Error** | 38 | 0.00617 | 0.000162 | | |
| **Corrected Total** | 47 | 0.0275 | | | |

Table S7. Factors affecting mean population growth rate at generation 4000. Standard least squares fit of mean growth rate of each population at generations 3875-4125 with all factors treated as fixed effects.

| **Source** | **DF** | **Sum of Squares** | **Mean Square** | **F Ratio** | **Prob > F** |
|---|---|---|---|---|---|
| **Model** | 26 | 0.425 | 0.0164 | 37.57 | <0.0001 |
| assay temperature | 1 | 0.271 | | 622.30 | <0.0001 |
| genotype | 2 | 0.0155 | | 1.39 | 0.270 |
| population[genotype]&Random | 21 | 0.128 | | 13.95 | <0.0001 |
| genotype*assay temperature | 2 | 0.00599 | | 6.87 | 0.0011 |
| **Error** | 826 | 0.360 | 0.000436 | | |
| **Corrected Total** | 852 | 0.785 | | | |

Table S8. Factors affecting mean population growth rate per plate at generation 4000. Standard least squares fit of mean population growth rate per plate between generations 3875-4125 with population nested within genotype treated as a random effect.

| **Source** | **DF** | **Sum of Squares** | **Mean Square** | **F Ratio** | **Prob > F** |
|---|---|---|---|---|---|
| **Model** | 26 | 0.131 | 0.00503 | 52.77 | <0.0001 |
| assay temperature | 1 | 0.0816 | | 855.65 | <0.0001 |
| genotype | 2 | 0.0410 | | 81.66 | <0.0001 |
| population[genotype]&Random | 21 | 0.00532 | | 2.65 | 0.0002 |
| genotype*assay temperature | 2 | 0.00189 | | 9.90 | <0.0001 |
| **Error** | 283 | 0.0270 | 0.000095 | | |
| **Corrected Total** | 309 | 0.158 | | | |

Table S9. Factors affecting mean population growth rate per plate at generation 0. Standard least squares fit of mean population growth rate per plate between generations 0-125 with population nested within genotype treated as a random effect.

| **Model #** | **Additional terms** | **AICc** | **Evidence ratio** |
|---|---|---|---|
| 1 | none | -23731.5 | > 2.00E+60 |
| 2 | genotype | -24389.52 | > 2.00E+60 |
| 3 | temperature | -28431.22 | > 2.00E+60 |
| 4 | genotype + temperature | -30067.54 | 1.97E+60 |
| 5 | genotype*temperature | -30123.63 | 1.30E+48 |
| 6 | temperature*generations | -28564.98 | > 2.00E+60 |
| 7 | genotype*generations | -24402.66 | > 2.00E+60 |
| 8 | temperature*generation + genotype | -30248.79 | 8.66E+20 |
| 9 | genotype*generations + temperature | -30109.36 | 1.64E+51 |
| 10 | genotype*temperature*generations | -30345.21 | n.a. |

Table S10. Fits of the generalized additive mixed model with various combinations of the terms genotype, temperature, and generation included. Model 1 includes only generations and growth rate and is coded in R using the gam function. Additional terms were added to the model (*gam(growthrate~s(generations)+(additional terms and all lower order combinations), data=dat)*) and the AICc was calculated for each fit. The evidence ratio was calculated by using the difference between AICc of each model the lowest AICc recorded for model 10 which included the three-way interaction and all lower order interactions (ER = e^(0.5*$\Delta AIC$)). The ER indicates the extent to which the data favors model 10 over each of the other models revealing the significance of the three-way interaction term used in this model.

| **Term** | **Scaled Estimate** | **Std Error** | **T ratio** | **Prob>\|t\|** |
|---|---|---|---|---|
| Intercept | 0.1107 | 0.000252 | 438.79 | 0.0000 |
| generations | 0.0135 | 0.000466 | 28.9 | <0.0001 |
| 24ºC | -0.0226 | 0.000252 | -89.6 | 0 |
| 37ºC | 0.0226 | 0.000252 | 89.6 | 0 |
| Parent (A) | -0.0141 | 0.000354 | -39.9 | 1.46e-306 |
| Parent (B) | 0.00665 | 0.000358 | 18.6 | 8.68e-75 |
| Offspring (AxB) | 0.00749 | 0.000358 | 20.9 | 2.13e-93 |
| Generations*24ºC | -0.00424 | 0.000466 | -9.09 | 1.36e-19 |
| Generations*37ºC | 0.00424 | 0.000466 | 9.09 | 1.36e-19 |
| Generations*Parent (A) | 0.0036 | 0.000653 | 5.51 | 3.72e-8 |
| Generations*Parent (B) | -0.000881 | 0.000662 | -1.33 | 0.183 |
| Generations*Offspring (AxB) | -0.00272 | 0.000663 | -4.1 | 0.0000417 |
| 24ºC*Parent (A) | -0.00219 | 0.000354 | -6.17 | 7.37e-10 |
| 24ºC*Parent (B) | 0.0018 | 0.000358 | 5.03 | 5.09e-7 |
| 24ºC*Offspring (AxB) | 0.000387 | 0.000358 | 1.08 | 0.28 |
| 37ºC*Parent (A) | 0.00219 | 0.000354 | 6.17 | 7.37e-10 |
| 37ºC*Parent (B) | -0.0018 | 0.000358 | -5.03 | 5.09e-7 |
| 37ºC*Offspring (AxB) | -0.000387 | 0.000358 | -1.08 | 0.28 |
| 24ºC*Parent (A)*Generations | **-0.00138** | 0.000653 | -2.11 | 0.035 |
| 24ºC*Parent (B)*Generations | 0.00121 | 0.000662 | 1.83 | 0.0678 |
| 24ºC*Offspring (AxB)* Generations | 0.000169 | 0.000663 | 0.255 | 0.799 |
| 37ºC*Parent (A)*Generations | **0.00138** | 0.000653 | 2.11 | 0.035 |
| 37ºC*Parent (B)*Generations | -0.00121 | 0.000662 | -1.83 | 0.0678 |
| 37ºC*Offspring (AxB)* Generations | -0.000169 | 0.000663 | -0.255 | 0.799 |

Table S11. Scaled estimates of all terms including all pairwise and higher order interactions generated from a standard least square model of mean growth rate per plate assayed at the evolution temperature. The scaled estimates in **bold** show the slower growing parent by generation by each temperature. The significantly negative value of this estimate at 24ºC and positive value at 37ºC is indicative of the faster convergence amongst the genotypes at the higher temperature.

### Figures

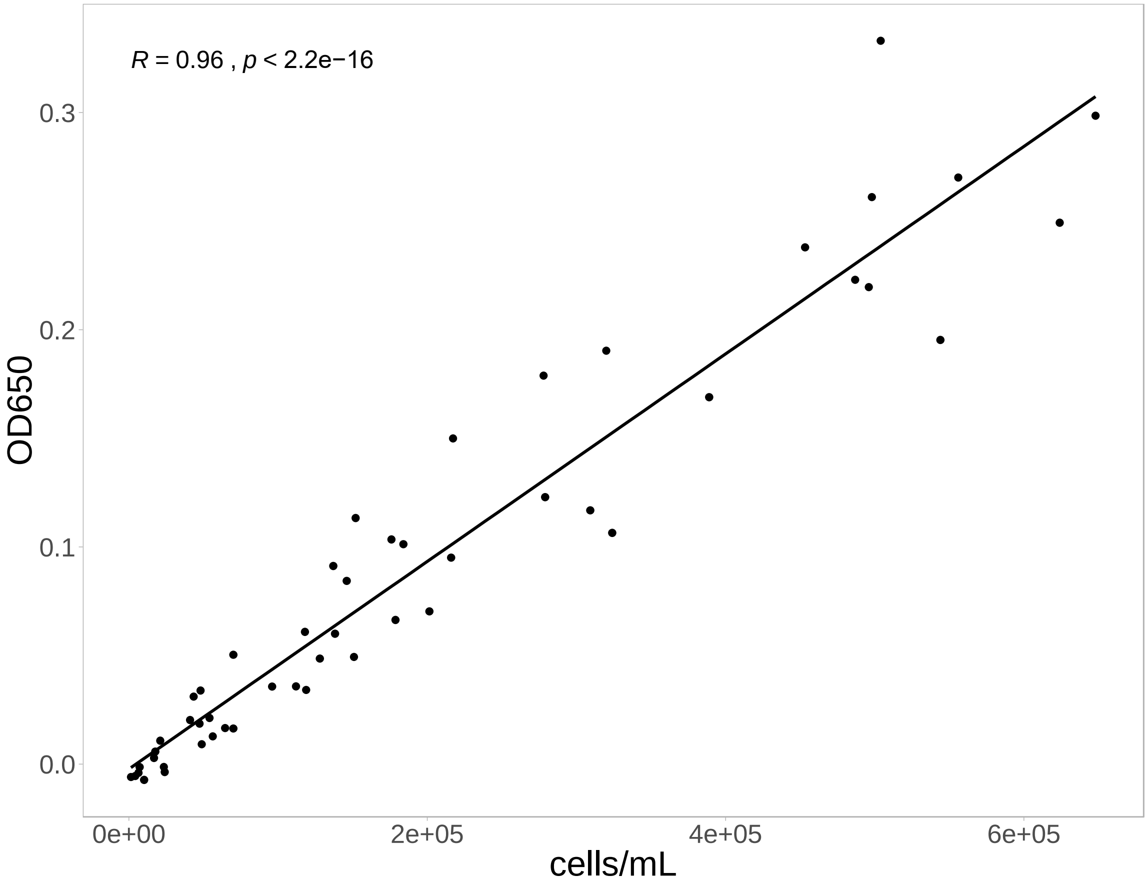


Figure S1. Correlation between OD650 and manual cell count. Each point shows the increase in OD650 attributable to cell growth and the manual cell count of a replicate population as it grows from low density to stationary phase. The black line shows the linear regression through this data. Pearson’s correlation coefficient is shown in the upper left corner.

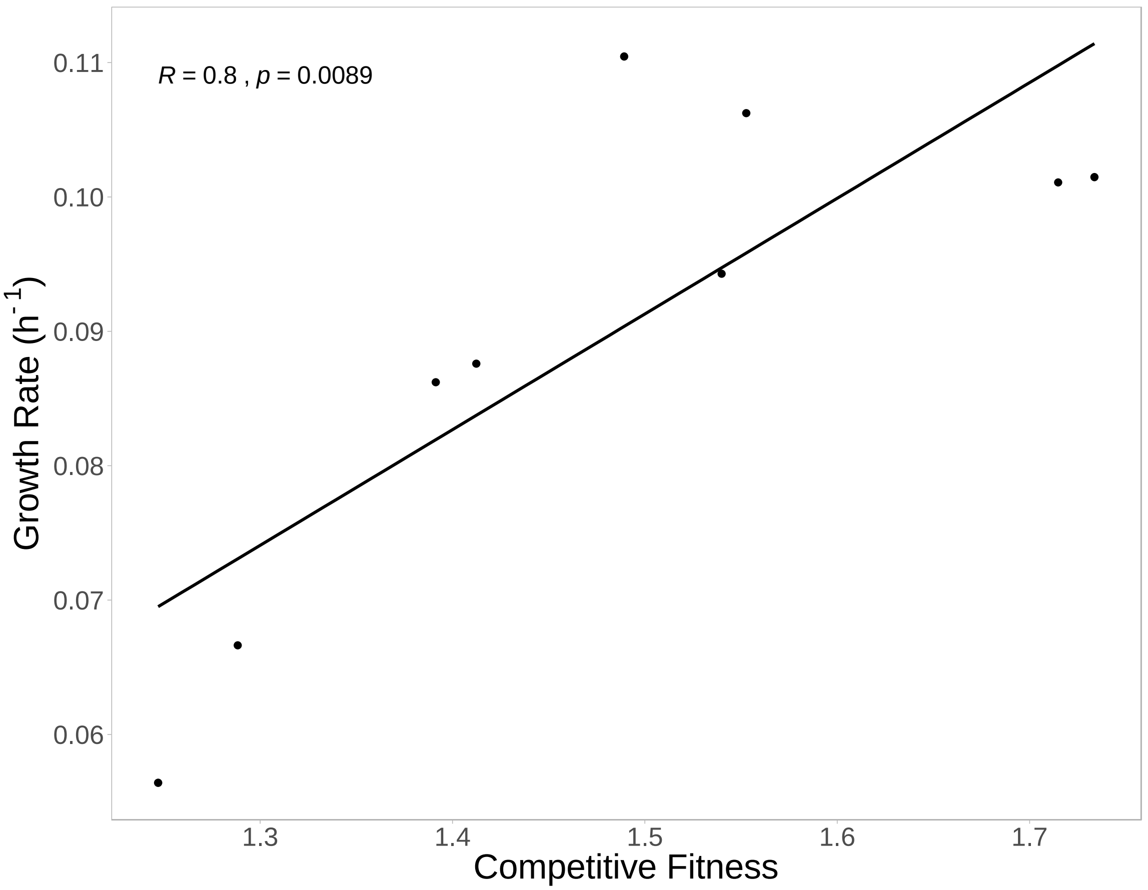


Figure S2. Correlation between growth rate and competitive fitness. Each point shows the growth rate and competitive fitness of one of nine populations for which competitive fitness was measured. The black line shows the linear regression through this data. Pearson's correlation coefficient is shown in the upper left corner.
